# Transient *DUX4* expression induces blastomere-like expression program that is marked by SLC34A2

**DOI:** 10.1101/2021.08.25.457357

**Authors:** Masahito Yoshihara, Ida Kirjanov, Sonja Nykänen, Joonas Sokka, Jere Weltner, Karolina Lundin, Lisa Gawriyski, Eeva-Mari Jouhilahti, Markku Varjosalo, Mari H. Tervaniemi, Timo Otonkoski, Ras Trokovic, Shintaro Katayama, Sanna Vuoristo, Juha Kere

**Author notes:** Correspondence (M.Y.), (S.V.), (J.K.). These authors contributed equally (Masahito Yoshihara, Ida Kirjanov).

## Abstract

DUX4 has recently been recognized as a key regulator in human embryonic genome activation (EGA). The exact role of DUX4 in human embryo is still elusive, partly due to the cytotoxicity of persistent *DUX4* expression in cellular models. We report here that a transient *DUX4* expression in human embryonic stem cells (hESCs) retains cell viability while inducing an EGA-like expression program in a subpopulation of the cells. These cells showed resemblance to 8-cell stage blastomeres and were thus named induced blastomere-like (iBM) cells. Trajectory inference from the single-cell RNA-seq data suggested that the expression profile of these cells progressed in a manner similar to the morula to blastocyst transition in human embryo. Finally, viable iBM cells could be enriched using an antibody against NaPi2b (SLC34A2), paving the way for further experimental approaches. The iBM cells can become a powerful tool to model transcriptional dynamics and regulation during early human embryogenesis.

## Introduction

DUX4, a double homeobox transcription factor, is a key regulator of human embryonic genome activation (EGA) by inducing cleavage stage-specific genes and repetitive elements (De Iaco et al., 2017; Hendrickson et al., 2017; Vuoristo et al., 2020; Whiddon et al., 2017). It is specifically and transiently expressed in the human cleavage-stage embryo (Töhönen et al., 2017). Activating mutations in the *DUX4* locus cause facio-scapulo-humeral muscular dystrophy (FSHD) (Tawil et al., 2014), where the ectopic expression of *DUX4* in skeletal muscles induces apoptosis via several pathways (Rickard et al., 2015; Shadle et al., 2017). *DUX4* overexpression in muscle cells leads to cell death both *in vitro* and *vivo* (Kowaljow et al., 2007; Wallace et al., 2011). However, transient *DUX4* expression in human myoblasts activates its target genes with little cytotoxicity by inducing histone variants H3.X and H3.Y, which contribute to the perdurance of DUX4 target gene expression with the open chromatin conformation (Resnick et al., 2019).

A rare cell population of mouse embryonic stem cells (mESCs) exhibit 2-cell-stage like signatures (Macfarlan et al., 2012), with the reduced expression of pluripotency markers such as *Pou5f1, Sox2*, and *Nanog*, and increased expression of targets of Dux, the mouse ortholog of human DUX4 (Rodriguez-Terrones et al., 2018). These 2-cell-like cells (2CLCs) spontaneously transit towards the pluripotent state under the culture conditions optimal for mESCs (Macfarlan et al., 2012). Recent studies have revealed that 2CLCs can be induced by *Dux* expression (De Iaco et al., 2017; Fu et al., 2019; Hendrickson et al., 2017; Yang et al., 2020). To the best of our knowledge, such 2CLCs have not been reported in human. These findings prompted us to examine whether human cells could be converted into an early embryonic-like state by transient *DUX4* expression that mimics its expression pattern during the early embryonic development.

## Results

### Transient DUX4 induction activates EGA genes in hESCs with little cytotoxicity

To test whether human ESCs (hESCs) can survive transient *DUX4* induction, we first examined the viability of doxycycline-inducible DUX4-TetOn hESCs after various durations of doxycycline exposure (15 min, 30 min, 1 h, and constant). While induction for 1 h or longer caused vast cell death, 15 min of induction resulted in a temporary decrease in growth rate, returning to a similar level with that of the cells without induction (**Figures 1A** and **S1A**). Moreover, only a small number of apoptotic cells were detected after 15 min of induction, at levels similar to cells without induction. Importantly, DUX4-positive cells were not positive for cleaved caspase-3, implying that transient *DUX4* expression did not induce apoptosis in hESCs (**Figure 1B**).

**Figure 1.**
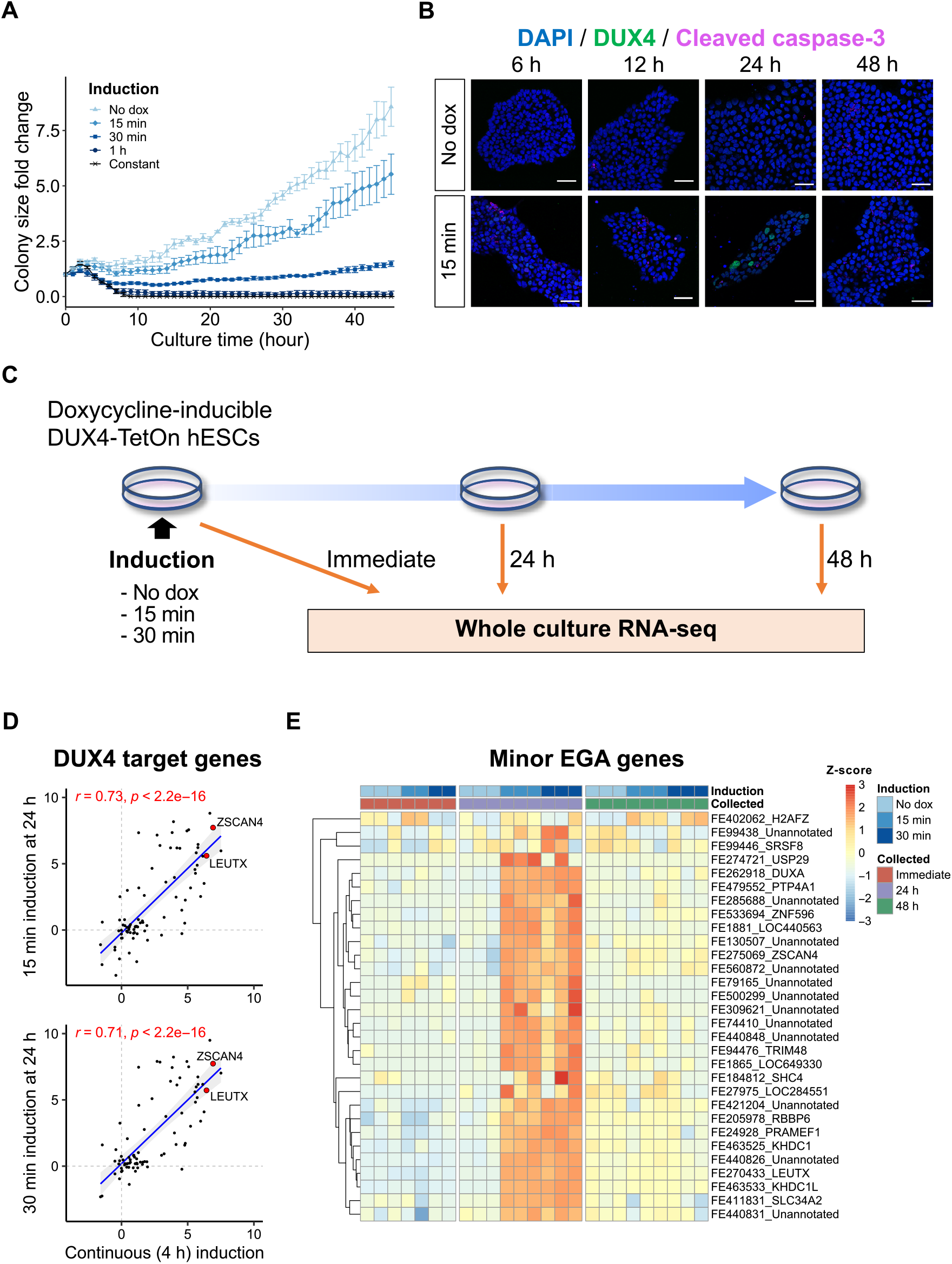
Transient *DUX4* induction activates EGA genes with little cellular toxicity. (A) Growth rate of DUX4-TetOn hESCs after varied times of doxycycline induction. Colony size fold change was calculated based on the starting time point (0 h). Data represents mean ± SEM (n=5). (B) Immunocytochemical detection of DUX4 and cleaved caspase-3 in DUX4-TetOn hESCs after 15 min induction (top) and without induction (bottom). DAPI (blue) was used as nuclear counterstain. Scale bars, 20µm. (C) Schematic representation of the whole culture RNA-seq on DUX4-TetOn hESCs with varied induction times. (D) Transcriptional changes of 80 DUX4 target genes expressed in early human embryo after continuous *DUX4* induction (x-axis) and transient *DUX4* induction (y-axis). Axes show the log_2_ fold expression changes over no *DUX4* induction. Spearman’s correlation coefficients (r) and *P*-values are indicated. (E) Heatmap showing the expression of minor EGA genes at TFE-level. A total of 30 expressed TFEs are shown. TFE, transcript far 5’ ends. See also Figure S1.

Next, to determine whether transient *DUX4* induction is sufficient to activate its target genes, DUX4-TetOn hESCs were exposed to doxycycline for 15 or 30 min and subjected to RNA sequencing (RNA-seq) (**Figure 1C**). Principal component analysis (PCA) demonstrated that cells collected at 24 h after either 15 or 30 min of induction were clearly separated from other samples along PC2 which was highly contributed by *LEUTX* and *ZSCAN4*, suggesting that short induction had the greatest effect at 24 h post-induction (**Figure S1B**). Notably, cells at 24 h after 15 and 30 min of induction showed highly similar expression profiles (r = 0.96, Spearman correlation) (**Figure S1C**). The expression changes of DUX4 target genes present in early embryo (Resnick et al., 2019) were similar between continuously (4 h) treated cells (Vuoristo et al., 2020), and pulsed cells at 24 h post-induction, suggesting that the 15 and 30 min pulse were sufficient to activate DUX4 target genes (**Figures 1D** and **S1D**). Furthermore, cleavage stage-specific repetitive elements such as MLT2A1, MLT2A2, and HERVL, activated by DUX4 (Geng et al., 2012; Hendrickson et al., 2017; Young et al., 2013), were significantly upregulated at 24 h after short induction (**Figure S1E**).

We previously investigated the dynamics of the human preimplantation transcriptome by Single-cell Tagged Reverse Transcriptase (STRT) RNA-seq quantifying the transcript far 5’ ends (TFEs), and identified 32 TFEs upregulated at the 4-cell stage as minor EGA genes (Töhönen et al., 2015), most of which should be regulated by DUX4 (De Iaco et al., 2017). We found that 30 of them were expressed in the *DUX4*-induced hESCs, and most of them showed the highest expression at 24 h after short induction (**Figure 1E**). Their expression was reduced at 48 h after induction, suggesting that the transient *DUX4* expression in hESCs recapitulated the transcriptional dynamics of cleavage stage-specific genes of early human embryos. These observations suggest that hESCs might be able to be converted into an early blastomere-like state with only a 15 min of *DUX4* induction.

### Transient DUX4 induction reprograms hESCs into an 8-cell-like transcriptional state

To examine whether early embryonic-like cells arise after transient *DUX4* induction, we performed time-series single-cell RNA-seq (scRNA-seq) on the DUX4-TetOn hESCs treated with doxycycline for 15 min (**Figure 2A**). We added two earlier time points before 24 h because the expression of DUX4 target genes might peak earlier, given their temporal expression in the embryo (De Iaco et al., 2017; Hendrickson et al., 2017). After filtering out low quality cells, 65,460 cells were retained for downstream analyses (**Table S1**). Dimensionality reduction by Uniform Manifold Approximation and Projection (UMAP) demonstrated that untreated (No dox) cells formed one main cluster, and a subset of *DUX4*-pulsed cells were separated from the main cluster (**Figure S2A**). The majority of *DUX4*-pulsed cells clustered together with the untreated cells, since induction did not affect the entire cell population. *DUX4* and its target genes were highly expressed in the rightmost cells along the first UMAP dimension (**Figures 2B** and **S2B**). Approximately 30–40% of cells expressed *DUX4* and its target genes at 6 h and 12 h, whereas only ∼5% of cells expressed them at 24 h (**Figure S2C**), in accordance with the small expression changes at 24 h in the whole-culture RNA-seq data (**Figure S1B**). EGA genes were specifically expressed in the rightmost cluster (**Figure 2C**).

**Figure 2.**
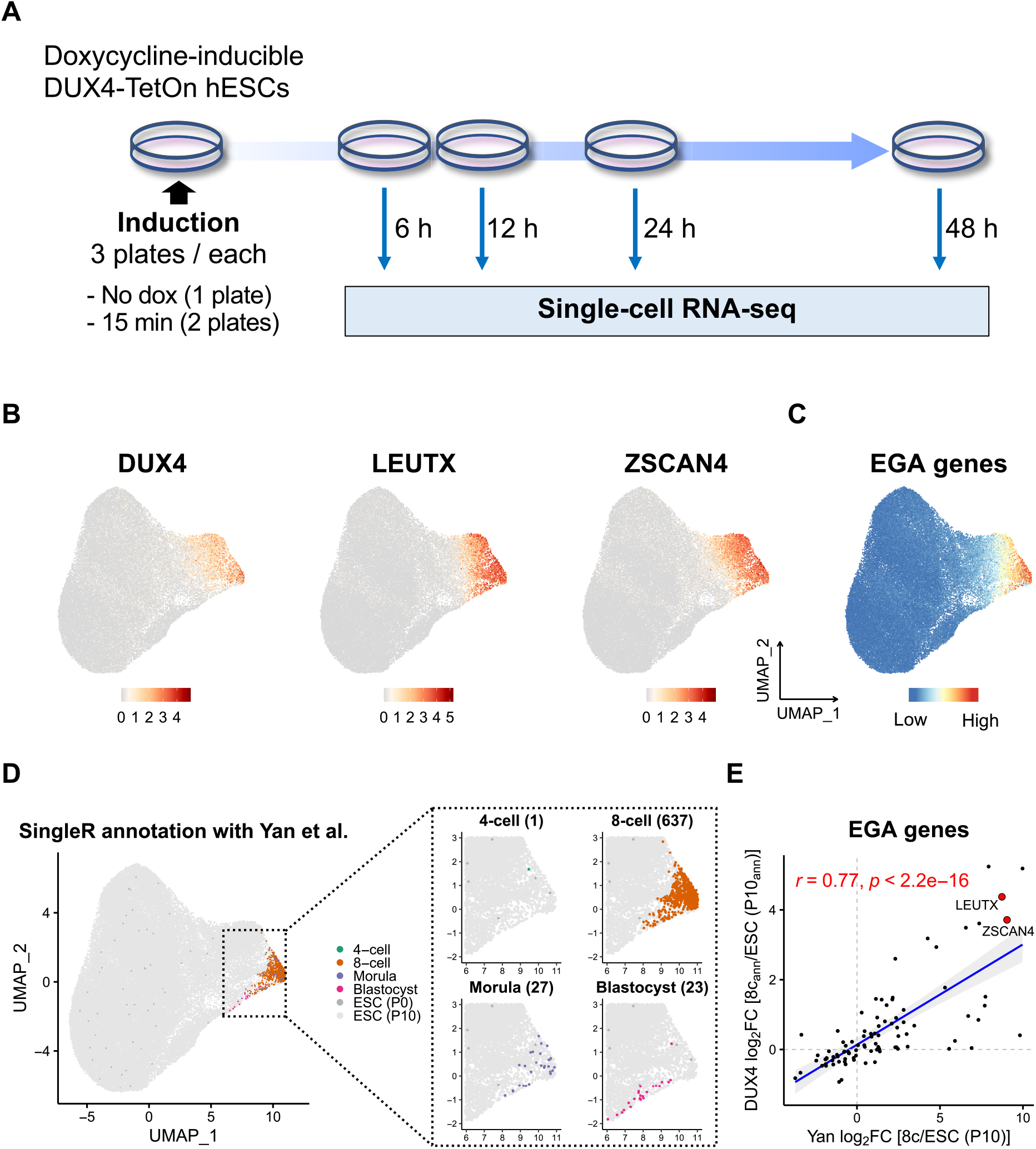
Time-series single-cell RNA-seq on *DUX4*-pulsed hESCs. (A) Schematic representation of the scRNA-seq experiments. (B) Expression of *DUX4, LEUTX*, and *ZSCAN4* projected onto the UMAP plot. (C) Expression score of EGA genes projected onto the UMAP plot. (D) Cell type annotation with human preimplantation embryos and hESCs using SingleR. The right four panels show the magnified plots of the cells annotated as early embryonic stage cells. Numbers in parentheses indicate the number of the annotated cells. P0, passage 0; P10, passage 10. (E) Transcriptional changes of 92 expressed EGA genes in actual 8-cell stage cells (x-axis) and *DUX4*-pulsed cells annotated as 8-cell stage cells (y-axis) compared with hESCs. Axes show the log_2_ fold expression changes over hESCs (P10; x-axis) or cells annotated as hESCs (P10; y-axis). 8c_ann_, cells annotated as 8-cell stage cells; ESC (P10_ann_), cells annotated as ESC (P10). See also Figure S2.

To address the similarity of these cells with actual early human embryonic cells, we annotated the cells against the scRNA-seq data of preimplantation embryos and hESCs (Yan et al., 2013) using SingleR (Aran et al., 2019). Strikingly, 637 cells were annotated as 8-cell stage cells (**Figure 2D**). Of these, 544 cells were collected at 12 h after induction. This indicates that 6.6% of the *DUX4*-pulsed cells (8,268 cells) were converted to a state that transcriptionally resembled 8-cell state 12 h after induction. Transcriptional changes of EGA genes in these 8-cell-like cells correlated highly with those in actual 8-cell stage cells (r = 0.77, Spearman correlation) (**Figure 2E**). Transcriptional changes of all the expressed genes were less correlated (r = 0.5; **Figure S2D** left), most likely reflecting the remaining maternal transcripts in 8-cell stage cells that were not activated by *DUX4* induction. In support of this, among the genes highly expressed in 8-cell stage cells (log_2_ FC > 5 over ESCs), genes activated by *DUX4* induction sharply increased from 2-to 8-cell stages, while genes not activated by *DUX4* induction were highly expressed in oocytes and zygotes (**Figure S2D** right). Of note, cells annotated as 4-cell, 8-cell, or morula were predominant at 12 h, whereas cells annotated as blastocyst were predominant at 24 h and 48 h (**Figure S2E**). This suggested that the transient *DUX4* pulsed cells might recapitulate aspects of the transcriptional dynamics of early human embryo.

### Cell-state transition dynamics after transient DUX4 induction

To further characterize the *DUX4*-pulsed cells, we assigned them to 6 clusters by unsupervised clustering. Based on the proportion of the collected time points and cell type annotations, we named the clusters as follows: Non-induced, Intermediate, induced blastomere-like (iBM), and Late 1, 2, and 3 (**Figure 3A**). The Intermediate cell cluster consisted primarily of 6 h sample cells, the iBM cluster of 12 h sample cells, and the Late clusters of 24 h and 48 h sample cells. Hierarchical clustering of these 6 clusters demonstrated that the iBM cluster showed a unique expression profile, whereas the Late 2 and 3 clusters shared similar profile with the Non-induced cluster (**Figure S3A**). The majority of the iBM cluster-specific genes (**Table S2**) were most highly expressed at 8-cell stage and downregulated in blastocyst (**Figure S3B**). The Intermediate and Late 1 clusters moderately expressed these genes with different patterns. Although none of these genes were expressed in the Non-induced cluster, *CCNA1* and *ALPG* were expressed in the Late 2 and 3 clusters. LEUTX target genes (**Table S3**) (Gawriyski et al., unpublished) were expressed higher in the Late 2 and 3 clusters than in the Non-induced cluster (**Figure S3C**). As *LEUTX* expression peaked in the iBM cluster (**Figure S3B**), the Late 2 and 3 clusters were likely derived from the iBM cells, distinguishing them from the Non-induced cluster.

**Figure 3.**
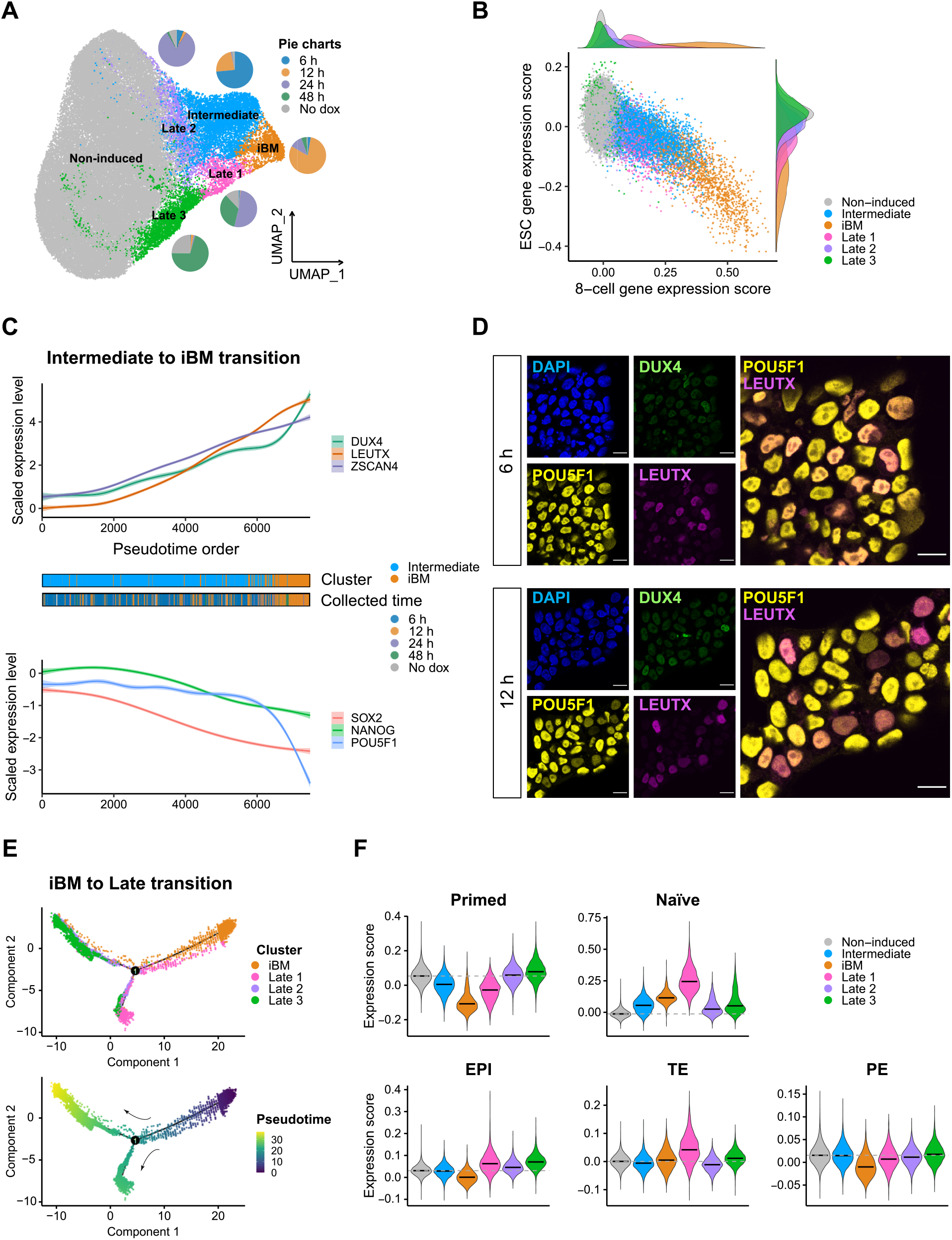
Clustering and trajectory analysis on *DUX4*-pulsed hESCs. (A) Clustering analysis of *DUX4*-pulsed hESCs with assigned cluster names. Pie charts represent the proportion of collected time points across the clusters. (B) Expression scores of 8-cell (x-axis) and ESC (y-axis) of each cell, colored by clusters. Distribution of cells of each cluster is shown at the top and right. (C) Expression changes of *DUX4* and its target genes (top) and pluripotency marker genes (bottom) of single cells from the Intermediate and the iBM clusters along the pseudotime. Middle panels show the cluster (above) and collected time point (below) of each single cell. (D) Immunocytochemical detection of DUX4, LEUTX, and POU5F1 in *DUX4*-pulsed hESCs at 6 h (top) and 12 h (bottom). DAPI (blue) was used as nuclear counterstain. Scale bars, 20 µm. (E) Trajectory reconstruction of single cells from the iBM and Late clusters: pre-branch (before bifurcation), Late 1 fate, Late 2–3 fate (after bifurcation). Top panel is colored by cluster and bottom one is colored by pseudotime. (F) Expression scores of primed and naïve PSC marker genes (top) and epiblast (EPI), trophectoderm (TE), and primitive endoderm (PE) (bottom) in each cluster. The horizontal black bars in violin plots indicate the median score per cluster, and the horizontal gray dotted line indicates the median score in the Non-induced cluster. See also Figure S3.

To estimate the reprogramming state changes from hESCs to iBM cells, we calculated the 8-cell and ESC gene expression scores in each cell, based on our scRNA-seq data on 8-cell stage cells and hESCs (**Table S3**) (Jouhilahti et al., 2016). As expected, cells in the Non-induced cluster showed the highest ESC score with a low 8-cell score, whereas cells in the iBM cluster showed the lowest ESC score with a high 8-cell score (**Figure 3B**). Cells in the Intermediate cluster located between these, suggesting that these cells were in the midst of the reprogramming process. Cells in the Late clusters showed higher ESC scores with lower 8-cell scores, suggesting that the iBM cells continued to develop toward the state of ESCs.

To dissect the reprogramming process to the iBM cells via Intermediate cells, we performed a pseudotime trajectory analysis on the 7,478 cells from these two clusters. The pseudotime order was consistent with the actual collected time points, with cells collected at 6 h being earlier and 12 h later (**Figure 3C**). The 8-cell and ESC gene expression scores showed an inverse changing pattern along the pseudotime (**Figure S3D**). Expression of *DUX4* and its targets increased along the pseudotime, whereas that of pluripotency marker genes such as *SOX2, NANOG*, and *POU5F1* decreased, although *POU5F1* did not significantly (**Figures 3C** and **S3E**; **Table S4**). These pluripotency marker genes are lowly expressed in cleavage-stage human embryos (Töhönen et al., 2015). DUX4 and LEUTX proteins were not detected in the untreated cells but were positive at 6 h and 12 h (**Figures 3D** and **S3F**). LEUTX-positive cells showed reduced POU5F1 staining, especially at 12 h (**Figure 3D**), as observed in mouse 2CLCs (Hendrickson et al., 2017; Macfarlan et al., 2012; Rodriguez-Terrones et al., 2018).

Next, to monitor the expression changes after the iBM stage, a pseudotime analysis was conducted on the 8,101 cells from the iBM and three Late clusters. Cells from the iBM cluster bifurcated into two diverse branches, Late 1 and Late 2–3 (**Figure 3E**). Interestingly, most of the naïve pluripotent stem cell (PSC) markers (Liu et al., 2020) were upregulated along the pseudotime progression in the Late 1 lineage, whereas primed PSC markers were highly expressed in the Late 2–3 lineage (**Figures 3F** and **S3G**). Since naïve PSCs have been reported to have a similar expression profile as preimplantation epiblast cells (Liu et al., 2017; Stirparo et al., 2018), we calculated the gene expression scores of epiblast (EPI), trophectoderm (TE), and primitive endoderm (PE) of all the cells in each cluster (Petropoulos et al., 2016). Intriguingly, cells in the Late 1 cluster showed the highest EPI score as well as TE score (**Figures 3F**). TE markers such as the S100 calcium binding protein coding genes, *DNMT3L*, and *GATA3* (Assou et al., 2012) were specifically expressed in the Late 1 cluster (**Figure S3H**). These genes were not highly expressed in the iBM cluster, implying that their expression was the result of a secondary effect after *DUX4* induction. These results suggest that the iBM cells might be primed to initiate the developmental progression towards the first embryonic lineages, inner cell mass and TE (Petropoulos et al., 2016).

### Viable iBM cells can be enriched with an anti-NaPi2b antibody

Finally, we aimed to enrich the iBM cells using an antibody against a cell surface antigen specifically expressed in the iBM cluster. Among the iBM-specific marker genes (**Table S2**), we discovered *SLC34A2*, encoding the sodium-dependent phosphate transporter NaPi2b, as the most specific cell surface marker gene (**Figure 4A**). *SLC34A2* is also one of the DUX4 target genes (Hendrickson et al., 2017) as well as EGA genes that is highly upregulated at the 4- and 8-cell stages (**Figure 1E**) (Töhönen et al., 2015), but rarely expressed in hESCs (**Figure 4B**). Mouse 2CLCs also highly express *Slc34a2* (Hendrickson et al., 2017). Similar to other DUX4 target genes, *SLC34A2* was highly expressed at 6 h and 12 h after induction (**Figure S4A**). We confirmed its expression at protein level with the anti-NaPi2b monoclonal antibody MX35 which recognizes its extracellular domain (Yin et al., 2008). MX35 specifically stained the cell surface of *DUX4*-pulsed hESCs, already at 6 h after induction (**Figures 4C** and **S4B**).

**Figure 4.**
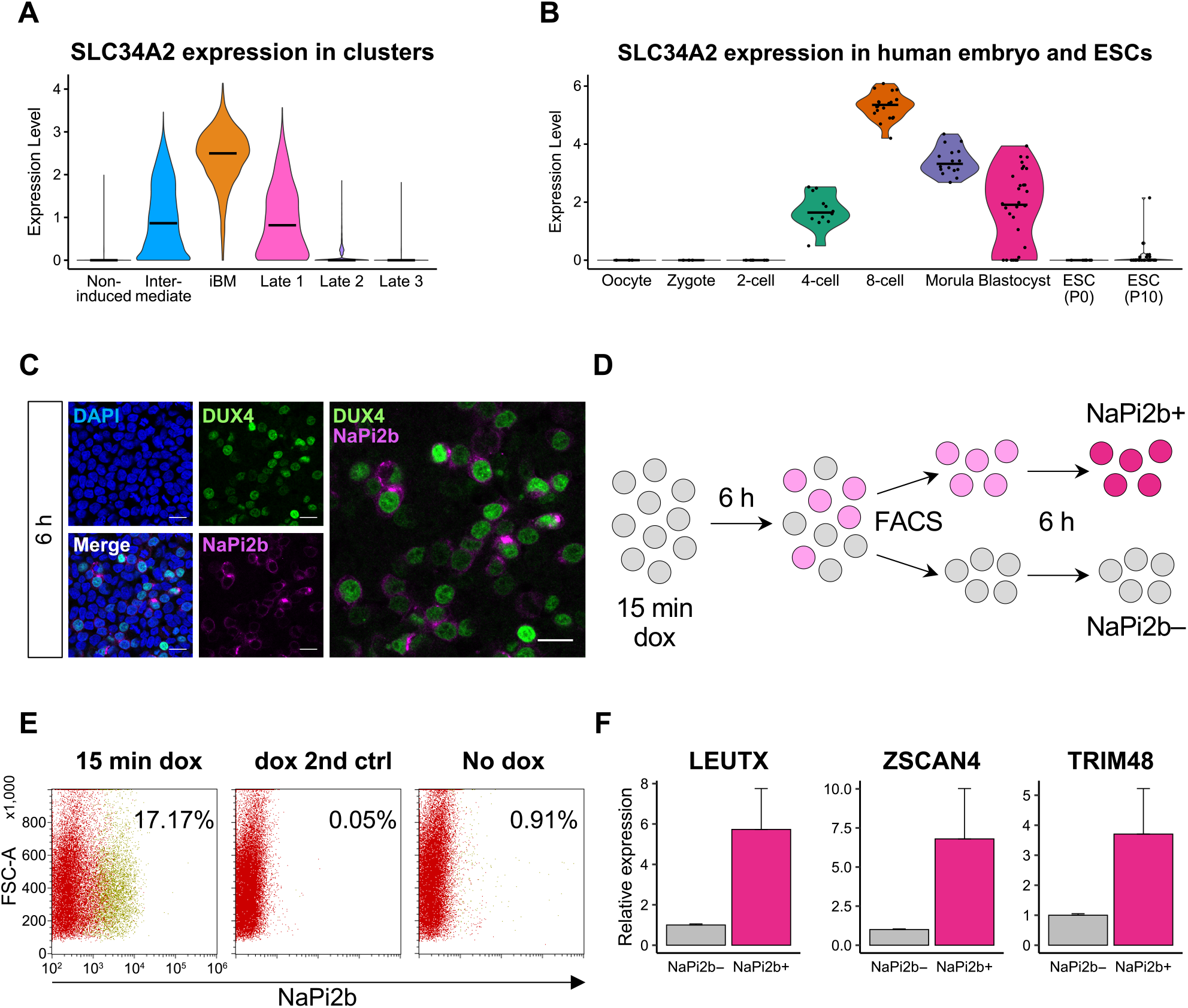
Enrichment of viable iBM cells with an anti-NaPi2b antibody. (A) *SLC34A2* expression in *DUX4*-pulsed hESCs in each cluster. Expression levels are shown as log normalized UMI counts. (B) *SLC34A2* expression in human preimplantation embryo and ESCs (Yan et al., 2013). Expression levels are shown as log FPKM values. The horizontal bars in each violin plot indicate median expression level per cluster (A) or stage (B). (C) Immunocytochemical detection of DUX4 and NaPi2b in *DUX4*-pulsed hESCs at 6 h. DAPI (blue) was used as nuclear counterstain. Scale bars, 20 µm. (D) Schematic illustration of the cell sorting procedure. (E) Flow cytometric analysis showing the proportion of NaPi2b-positive cells (yellow dots). Representative data from two independent experiments are shown. FSC, forward scatter; 2nd ctrl, secondary antibody control. (F) DUX4 target gene expression in NaPi2b+/- cells cultured for 6 h after cell sorting. Error bars represent the SEM of two independent experiments. See also Figure S4.

We then attempted to enrich NaPi2b-positive cells by fluorescence-activated cell sorting (FACS) at 6 h after induction and cultured them for a further 6 h toward the fully reprogrammed state (**Figure 4D**), given that iBM cells were most enriched at 12 h after induction. The proportion of NaPi2b-positive cells in *DUX4*-pulsed hESCs was ∼15% (of two independent experiments; **Figures 4E** and **S4C**). After 6 h of culture, NaPi2b-positive cells expressed higher levels of DUX4 target genes than NaPi2b-negative cells (**Figure 4F**). Moreover, Annexin V staining showed that apoptotic rate between NaPi2b+/- cells was comparable (**Figure S4D**), even though dead cells could be observed among the NaPi2b-positive cells. These observations indicate that viable iBM cells for further cell culture could be enriched by FACS using the MX35 antibody.

## Discussion

We describe here the transcriptional reprogramming of primed hESCs into blastomere-like (iBM) cells by a transient *DUX4* induction. The iBM cells showed upregulation of EGA genes and downregulation of pluripotency factors. These cells could be marked and sorted out live with an antibody against NaPi2b.

Although a number of studies have investigated DUX4-mediated cytotoxicity (Bosnakovski et al., 2008; Geng et al., 2012; Rickard et al., 2015; Shadle et al., 2017; Wallace et al., 2011), its mechanism is not fully understood yet. As DUX4 is a key regulator of human EGA, it still remains unclear whether its toxic effect is avoided in embryos by the brief expression seen *in vivo* or by other mechanisms. The balance between cytotoxicity and cell survival after *DUX4* induction deserves further studies.

Our transcriptomic comparison with cleavage-stage embryos showed that EGA genes were efficiently activated by a transient *DUX4* induction, although many oocyte specific genes were absent. The relevance of these oocyte specific factors for modeling early embryo behavior with stem cells remains to be determined. The pluripotency markers *NANOG* and *SOX2* were significantly downregulated in the iBM cells. They showed reduced level of POU5F1 protein, although its expression was not significantly affected at transcriptional level. These pluripotency markers are also decreased in mouse 2CLCs (Hendrickson et al., 2017; Macfarlan et al., 2012; Rodriguez-Terrones et al., 2018). In accordance with the iBM cells, *Sox2* transcript level is downregulated in mouse 2CLCs. Although *Pou5f1* transcript is not affected, it is downregulated on a protein level (Fu et al., 2020; Rodriguez-Terrones et al., 2018). Even though mouse 2CLCs and human iBM cells share key features, the direct comparison of these cell types remains challenging due to the evolutionary divergence of Dux and DUX4 targets between the species (Whiddon et al., 2017). For example, of the DUX4 target genes in human cells, some of the human PRD-like factors such as *LEUTX*, have been replaced by the mouse *Obox* factors (Royall et al., 2018). *KLF17* has been functionally replaced by *Klf2* in mouse embryos (Stirparo et al., 2018).

Markers of naïve PSCs and trophectoderm were upregulated in a subpopulation of the cells, most likely emerging from the iBM cells. Intriguingly, recent papers have described the enhanced developmental capacity of naïve PSCs to produce extraembryonic cell types (Guo et al., 2021; Io et al., 2021; Linneberg-Agerholm et al., 2019; Yanagida et al., 2021). Therefore, DUX4 likely enhances the establishment of the late morula to early blastocyst differentiation capacity of embryos that may be mimicked in naïve PSCs.

Finally, the iBM cells were marked by the expression of *SLC34A2*, encoding NaPi2b. Importantly, this provides a way for detecting *DUX4*-induced cells in human PSC cultures. As the marker is extracellular, it does not require transgenic reporter constructs, like MERVL or *Zscan4* promoter driven fluorescent proteins (Macfarlan et al., 2012; Rodriguez-Terrones et al., 2018). We envision that MX35 may be of use for developing new ways of inducing human blastomere-like cells, without the need for transgenic *DUX4* expression, and for more stable ways of culturing human blastomere-like cells, as was recently described for mouse (Shen et al., 2021).

In conclusion, transient *DUX4* pulsing can be used to induce human blastomere-like cells, which can be enriched by an anti-NaPi2b antibody. These approaches will potentially allow the generation of improved cell models for the early human embryogenesis without the need for human embryos.

## Methods

### Cell culture

DUX4-TetOn hESCs (Vuoristo et al., 2020) were maintained on hESC-qualified Geltrex (Thermo Fisher Scientific) coated tissue culture dishes in Essential 8 culture medium (Thermo Fisher Scientific) in 5% CO_2_ at 37°C. The cells were passaged every three to five days after a 3 min incubation with 0.5 mM EDTA (Thermo Fisher Scientific). For the cell growth assays, the cells were imaged with the IncuCyte S3 analysis system (Sartorius).

### Doxycycline pulsing on DUX4-TetOn hESCs

DUX4-TetOn hESCs were incubated with 1 µg/ml of doxycycline in Essential 8 culture medium in 5% CO_2_ incubator at 37°C for varied times as indicated. After the doxycycline induction, the DUX4-TetOn hESCs were washed three times with Essential 8 culture medium and incubated thereafter in Essential 8 medium for the indicated times.

### RNA extraction and quantitative real-time (qRT)-PCR

Total RNA was isolated using NucleoSpin RNA kit (Macherey Nagel) according to the manufacturer’s protocol. For cDNA synthesis, 500 ng of total RNA was reverse-transcribed by MMLV-RTase (Promega) with oligo dT priming. The resulting cDNA was used as a template for qRT-PCR using 5× HOT FIREPol qPCR Mix (Solis BioDyne) on the LightCycler 96 System (Roche). Relative expression values were calculated with the 2^-ΔΔCt^ method (Livak and Schmittgen, 2001), using cyclophilin G (*PPIG*) as an internal control, normalized against the NaPi2b-negative cells.

### STRT whole culture RNA-seq library preparation and sequencing

Doxycycline-induced and control DUX4-TetOn hESCs were collected for STRT whole culture RNA-seq immediately, 24 h, and 48 h after 15 min, 30 min of doxycycline, or without doxycycline treatment. We used 20 ng of RNA to generate a 48-plex RNA-seq library using a modified STRT method with unique molecular identifiers (UMIs) (Islam et al., 2011; Islam et al., 2014). Briefly, RNA samples were placed in a 48-well plate, and a universal primer, template-switching oligo-nucleotides, and a well-specific 6 bp barcode sequence (for sample identification) were added to each well (Katayama et al., 2013; Krjutškov et al., 2016). We pooled the synthesized cDNAs into one library, performed fragmentation to 200–400 bp using an M220 Focused-ultrasonicator (Covaris), captured the 5’-prime fragments, added an adapter, and amplified the targets by PCR. The RNA-seq library was sequenced with Illumina NextSeq 500 System, High Output mode (75 cycles).

### scRNA-seq library preparation and sequencing

DUX4-TetOn hESCs were seeded into three plates at each experiment, two doxycycline-treated and one untreated, and then collected at 6 h, 12 h, 24 h, and 48 h after treatment. The cells were washed once with PBS and incubated with TrypLE Express (Thermo Fisher Scientific) for 4 min. TrypLE was diluted with Essential 8 medium and the cell suspensions were filtered through 40 µm Cell Strainers. Cell suspensions were centrifuged at 400 rcf for 8 min. Cell pellets were resuspended each in 100 µl of Dead Cell Removal Kit microbeads (Miltenyi Biotec) and incubated at room temperature for 15 min. After incubation, each cell-microbead suspension was gently resuspended to 800 µl of freshly prepared 1× Binding buffer. Cell suspensions were pipetted to magnetic MS columns (Miltenyi Biotec) 500 µl at a time and let flow through. The columns were washed three times with 1× Binding buffer. The cell suspensions were centrifuged at 400 rcf for 5 min, and the pellets were resuspended each in 400 µl of 10x Genomics sample buffer. The cells were counted and the volumes were adjusted to approximately 1,200 cells/µl of suspension. The samples were kept on ice prior to analysis of cell quality and number, and preparation of the scRNA sequencing libraries. Approximately 94% of the nucleated cells were alive. The libraries were prepared using Chromium Next GEM Single Cell 3’ Gene Expression version 3.1 chemistry and sequencing was performed using Illumina NovaSeq 6000 system at the Institute for Molecular Medicine Finland (FIMM) Single-Cell Analytics unit.

### Immunocytochemical staining of DUX4-TetOn hESCs

Cells were fixed on Ibidi 8-well *µ* slides with 3.8% paraformaldehyde at room temperature for 15 min and washed three times with PBS. For the nuclear epitopes the cells were permeabilized using 0.5% Triton X-100-PBS at room temperature for 7 min. The cells were washed once with PBS and unspecific binding of antibodies was blocked by Ultravision Protein Block solution (Thermo Fisher Scientific) by a 10 min incubation at room temperature. Primary antibodies were diluted in washing buffer (0.1% Tween20-PBS), and incubated at 4°C overnight. Excess primary antibody solutions were removed and the cells were washed three times with washing buffer. The secondary antibodies were diluted 1:1000 in washing buffer and incubated at room temperature for 30 min. The samples were washed three times with washing buffer and nuclei were counterstained with DAPI, diluted 1:1000 in washing buffer. The samples were washed once and kept in PBS for imaging.

### Confocal microscopy and image analysis

Images were captured with a Leica TCS SP8 confocal laser scanning microscope (Leica Microsystems, Mannheim, Germany) using Leica HC PL APO CS2 40×/1.10NA water objective and 1024×1024 scan format. For Annexin V stainings, the cells were imaged with a Leica TCS SP8 X confocal microscope with white laser. The images were captured with either 20× air objective or 63× oil objective using 1024×1024 scan format. The data were processed using Fiji (http://fiji.sc). The images were softened using Gaussian filter (radius = 1-pixel kernel).

### Fluorescence activated cell sorting (FACS) of DUX4-TetOn hESCs

The DUX4-TetOn hESCs were washed once with PBS and incubated with TrypLE Express for 4 min in 5 % CO_2_ incubator at 37°C. The TrypLE Express was diluted with cold FACS buffer (5% fetal bovine serum in PBS supplemented with 10 µM ROCK inhibitor Y-27632 (Selleckchem)) and the cell suspensions were let flow through 40 µm Cell Strainers. The cells were count and approximately 5×10^5^ cells were aliquoted per Eppendorf tube. From here on the cells were kept on ice. The cells were centrifuged at 4°C, 300 rcf for 5 min. The primary anti-NaPi2b antibody, mouse MX35, a kind gift from Dr. Gerd Ritter, Ludwig Institute for Cancer Research, was diluted 1:100 (final concentration 20 µg/ml) in FACS buffer. The cells were incubated for 1 h on ice for primary antibody staining (MX35). The samples were washed three times with FACS buffer by centrifugation as above. Secondary antibody Alexa 594-conjugated donkey anti-mouse (A-21203, Thermo Fisher Scientific), was diluted 1:1000 in FACS buffer and incubated with cells on ice for 30 min. The cells were washed three times as above. The cells were analyzed and separated using Sony SH800Z Cell Sorter (Sony Biotechnology), using 100 µm nozzle. Altogether 5×10^5^ cells were collected for follow-up culture. The cells were centrifuged at 4°C, 300 rcf for 5 min, resuspended in Essential 8 culture medium with 10 µM ROCK inhibitor, and cultured for 6 h in 5% CO_2_, at 37°C, prior to cell lysis for RNA isolation.

### Annexin V staining

The cells were washed with Annexin V binding buffer and incubated in Annexin V solution (1:100 in binding buffer) at room temperature in dark, for 5 min. The cells were washed with Annexin V Binding buffer, fixed with 2% paraformaldehyde at room temperature in dark, for 10 min. The cells were washed twice with PBS and imaged.

### STRT RNA-seq data processing

The sequenced STRT RNA-seq raw reads were processed as described elsewhere (Lauter et al., 2020). Briefly, raw base call (BCL) files were demultiplexed and converted to FASTQ files with Picard tools (v2.20.4; http://broadinstitute.github.io/picard/), and aligned to the human reference genome hg19, human ribosomal DNA unit (GenBank: U13369), and ERCC spike-ins (SRM 2374) with the GENCODE (v28) transcript annotation by HISAT2 (v2.1.0) (Kim et al., 2019). Potential PCR duplicates were flagged with Picard MarkDuplicates. For gene-based analysis, uniquely mapped reads within the 5’-UTR or 500 bp upstream of the protein-coding genes were counted using Subread featureCounts (v1.5.2) (Liao et al., 2014) with ‘−−ignoreDup’ option. The mapped reads were further assembled by StringTie (v1.3.3) (Pertea et al., 2015) and those reads within the first exons of the assembled transcripts (TFEs) were counted as previously described (Töhönen et al., 2015). Two samples collected immediately after induction were excluded due to a low number of mapped reads. PCA was performed using the top 500 most variable genes. The STRT RNA-seq data of continuous *DUX4* induction, treated by doxycycline for 4 h, was obtained from Vuoristo et al. (Vuoristo et al., 2020) and reprocessed as described above. The expression of transposable elements was quantified using TEtranscripts (v2.2.1) with ‘uniq’ mode (Jin et al., 2015). Differential expression analysis between doxycycline-induced and non-induced cells was performed with the R (v4.0.0) package DESeq2 (v1.30.0) (Love et al., 2014), and the expression values were normalized by the library size calculated with DESeq2. The list of EGA genes was retrieved from Töhönen et al. (Töhönen et al., 2015), and TFEs overlapped with these gene regions were analyzed further. The list of DUX4 target genes expressed in the cleavage-stage human embryo were retrieved from Resnick et al. (Resnick et al., 2019).

### scRNA-seq data processing

#### Data pre-processing and cell clustering

The raw BCL files were demultiplexed and converted to FASTQ files with Cell Ranger (10x Genomics, v3.1.0) mkfastq, and mapped against the customized human reference genome (GRCh38 with DUX4-IRES-EmGFP) with STAR (Dobin et al., 2013). The cellranger aggr pipeline was used to combine all the data to generate a gene-count matrix. The output count data were subsequently analyzed with the R package Seurat (v4.0.0) (Hao et al., 2021). Cells with 15,000–100,000 UMI counts, expressing over 3,500 genes and less than 15% mitochondrial counts were kept, resulting in 65,460 cells in total. These data were then processed using the NormalizeData and FindVariableFeatures (using 2,000 features) functions. Next, cell-cycle scores were calculated using the CellCycleScoring function, and data scaling was performed with the ScaleData function, regressing out the S and G2M scores. Principal component analysis (PCA) was performed on the scaled data using the RunPCA function, and cell clustering was performed using the FindNeighbours (using the top 10 PCs) and FindClusters (resolution = 0.6) functions. UMAP was implemented on the top 10 PCs with the RunUMAP function. Here, 10 clusters with lower UMAP_1 values (left clusters) showing similar expression profiles were mainly composed of No-dox cells and were assigned as the ‘Non-induced’ cluster. The dendrogram was generated with the BuildClusterTree function. To measure the expression of *DUX4*, we quantified the expression of DUX4-IRES-EmGFP to avoid problems of mapping to the D4Z4 repeat locus. Average expression level in each cluster was calculated with the AverageExpression function. The iBM cluster specific markers shown in Figure S3B were identified by the FindAllMarkers function and selected as pct.1 > 0.8, pct.2 < 0.5, and avg_logFC > 1.

#### Gene expression scoring and cell type annotation

Gene expression scores of each signature were calculated using the gene signature scoring function retrieved from Liu et al. (Liu et al., 2020). Briefly, the average expression values of the genes of interest were subtracted by the aggregated expression values of a set of randomly selected control genes at similar expression level. The list of EGA genes were obtained from Töhönen et al. (Töhönen et al., 2015), and that of signature genes of EPI, TE, PE, primed, and naïve were obtained from Liu et al. (Liu et al., 2020). The list of 8-cell and ESC genes were retrieved from our previous STRT RNA-seq data (Jouhilahti et al., 2016), where the top 121 and 119 differentially expressed genes based on the differential expression score by STRTprep (Krjutškov et al., 2016) were selected, respectively (**Table S3**). The list of 299 LEUTX-target genes were retrieved from the significantly upregulated genes in our unpublished STRT RNA-seq data on LEUTX-inducible hESCs (Gawriyski et al., unpublished). Cell type annotation was conducted with the R package SingleR (v1.4.1) (Aran et al., 2019), using the scRNA-seq data of human preimplantation embryos and ESCs (Yan et al., 2013) as the reference data.

#### Pseudotime trajectory analysis

Pseudotime trajectory analysis was performed using the R package Monocle (v2.18.0) (Qiu et al., 2017) for two groups of clusters: i) Intermediate and iBM clusters, ii) iBM and Late 1–3 clusters, respectively. Cluster marker genes (**Table S2**) identified by the FindAllMarkers function in Seurat were used for ordering the cells, and dimensionality was reduced using the DDRTree algorithm. A generalized additive model (GAM) was fitted to the scaled expression values calculated by Seurat and the pseudotime order of cells using the geom_smooth function of the R package ggplot2 (v3.3.3). Heatmaps were generated with the Monocle plot_pseudotime_heatmap and plot_genes_branched_heatmap functions.

## Acknowledgements

The authors thank Dr. Gerd Ritter (Ludwig Institute for Cancer Research, New York City, USA) for kindly providing anti-NaPi2b monoclonal antibody (MX35). The whole culture STRT RNA-seq and scRNA-seq were carried out at the Biomedicum Functional Genomics Unit (FuGU), University of Helsinki and FIMM Single-Cell Analytics unit supported by HiLIFE and Biocenter Finland, respectively. The flow cytometry analysis was performed at the HiLife Flow Cytometry Unit, University of Helsinki. Most of the computations for this work were performed on resources provided by the Swedish National Infrastructure for Computing (SNIC) through the Uppsala Multidisciplinary Center for Advanced Computational Science (UPPMAX) under project 2017/7-317. MY was supported by Scandinavia–Japan Sasakawa Foundation, Japan Eye Bank Association, Astellas Foundation for Research on Metabolic Disorders, and Japan Society for the Promotion of Science (JSPS) Overseas Research Fellowships. SV lab was supported by research project grants from Sigrid Jusélius Foundation and University of Helsinki. Work in the JK lab was supported by Jane & Aatos Erkko Foundation, Sigrid Jusélius Foundation, Swedish Research Council, and Föreningen Liv och Hälsa. JW was supported by Sigrid Jusélius Foundation and Päivikki and Sakari Sohlberg Foundation. SK was supported by Jane & Aatos Erkko Foundation.

## Author contributions

Conceptualization: M.Y., S.V., J.K.; Data curation: M.Y.; Formal Analysis: M.Y., J.S.; Funding acquisition: T.O., S.V., J.K.; Investigation: M.Y., I.K., S.N., J.S., K.L., L.G., E.-M.J., M.V., M.H.T., S.V.; Methodology: M.Y., J.W., R.T., S.V.; Project administration: J.K.; Resources: R.T., S.K., S.V.; Software: M.Y., S.K.; Supervision: T.O., R.T., S.K., S.V., J.K.; Validation: M.Y., I.K., S.N., S.V.; Visualization: M.Y., S.V.; Writing – original draft: M.Y., S.V.; Writing – review & editing: All authors

## Declaration of Interests

The authors declare no competing interests.

## Supplementary Tables

**Table S1. Quality metrics of the scRNA-seq data in each experiment, related to Figures 2 and 3**.

**Table S2. List of marker genes in each cluster, related to Figure 3**.

**Table S3. List of genes used for calculating the gene expression scores, related to Figure 3**.

**Table S4. List of differentially expressed genes along the pseudotime (q < 1e-100) from Intermediate to iBM transition, related to Figure 3**.

## Lead Contact

Further information and requests should be directed to the Lead Contact, Juha Kere.

## Data Availability

The whole culture RNA-seq and scRNA-seq data of DUX4-TetOn hESCs used in this study have been deposited in the ArrayExpress database at EMBL-EBI under the accession codes ‘E-MTAB-10569’ and ‘E-MTAB-10581’, respectively (*publicly available after acceptance*).

## Notes

### Competing Interest Statement

The authors have declared no competing interest.

