## Supplementary Figures for "Transient *DUX4* expression induces blastomere-like expression program that is marked by SLC34A2"

Fig. S1

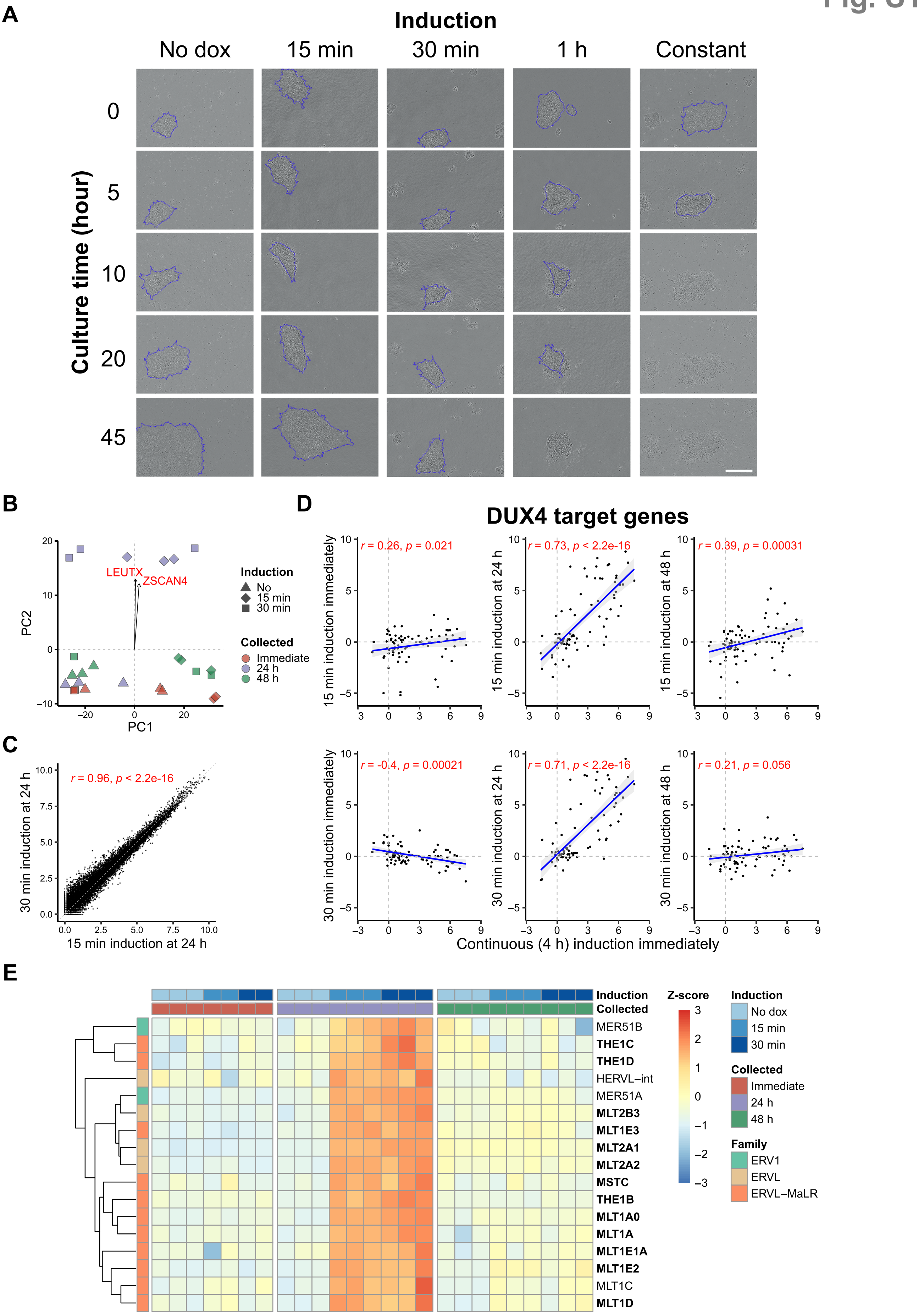

A
Induction
No dox
15 min
30 min
1 h
Constant
Culture time (hour)
0
5
10
20
45
B
PC2
PC1
LEUTX
ZSCAN4
Induction
No
15 min
30 min
Collected
Immediate
24 h
48 h
C
r = 0.96, p < 2.2e-16
30 min induction at 24 h
15 min induction at 24 h
D
DUX4 target genes
r = 0.26, p = 0.021
r = 0.73, p < 2.2e-16
r = 0.39, p = 0.00031
15 min induction immediately
15 min induction at 24 h
15 min induction at 48 h
r = -0.4, p = 0.00021
r = 0.71, p < 2.2e-16
r = 0.21, p = 0.056
30 min induction immediately
30 min induction at 24 h
30 min induction at 48 h
Continuous (4 h) induction immediately
E
Induction
Collected
MER51B
THE1C
THE1D
HERVL–int
MER51A
MLT2B3
MLT1E3
MLT2A1
MLT2A2
MSTC
THE1B
MLT1A0
MLT1A
MLT1E1A
MLT1E2
MLT1C
MLT1D
Z-score
Induction
No dox
15 min
30 min
Collected
Immediate
24 h
48 h
Family
ERV1
ERVL
ERVL–MaLR

**Figure S1. Effect of transient *DUX4* induction in hESCs, related to Figure 1**

(A) Bright field microscopy images of DUX4-TetOn hESCs after varied times of doxycycline induction. Living cells used for the measurement of colony size are surrounded by blue lines. Scale bars, 200 μm.

(B) Principal component analysis of the STRT whole culture RNA-seq data. Arrows show the variables of *LEUTX* and *ZSCAN4* on PC1 and PC2.

(C) Correlation of gene expression profiles of DUX4-TetOn hESCs at 24 h after 15 min (x-axis) and 30 min (y-axis) of induction. Expression levels are shown as log normalized counts.

(D) Transcriptional changes of 80 DUX4 target genes present in early human embryo after continuous *DUX4* induction (x-axis) and transient *DUX4* induction (y-axis). Axes show the $\log_2$ fold expression changes over no *DUX4* induction. Spearman's correlation coefficients (r) and *P*-values are indicated.

(E) Heatmap showing the expression of repetitive elements. Repetitive elements significantly upregulated at 24 h after 15 min and 30 min of induction are shown. Elements bound by DUX4 (Young et al., 2013) are shown in bold.

Fig. S2

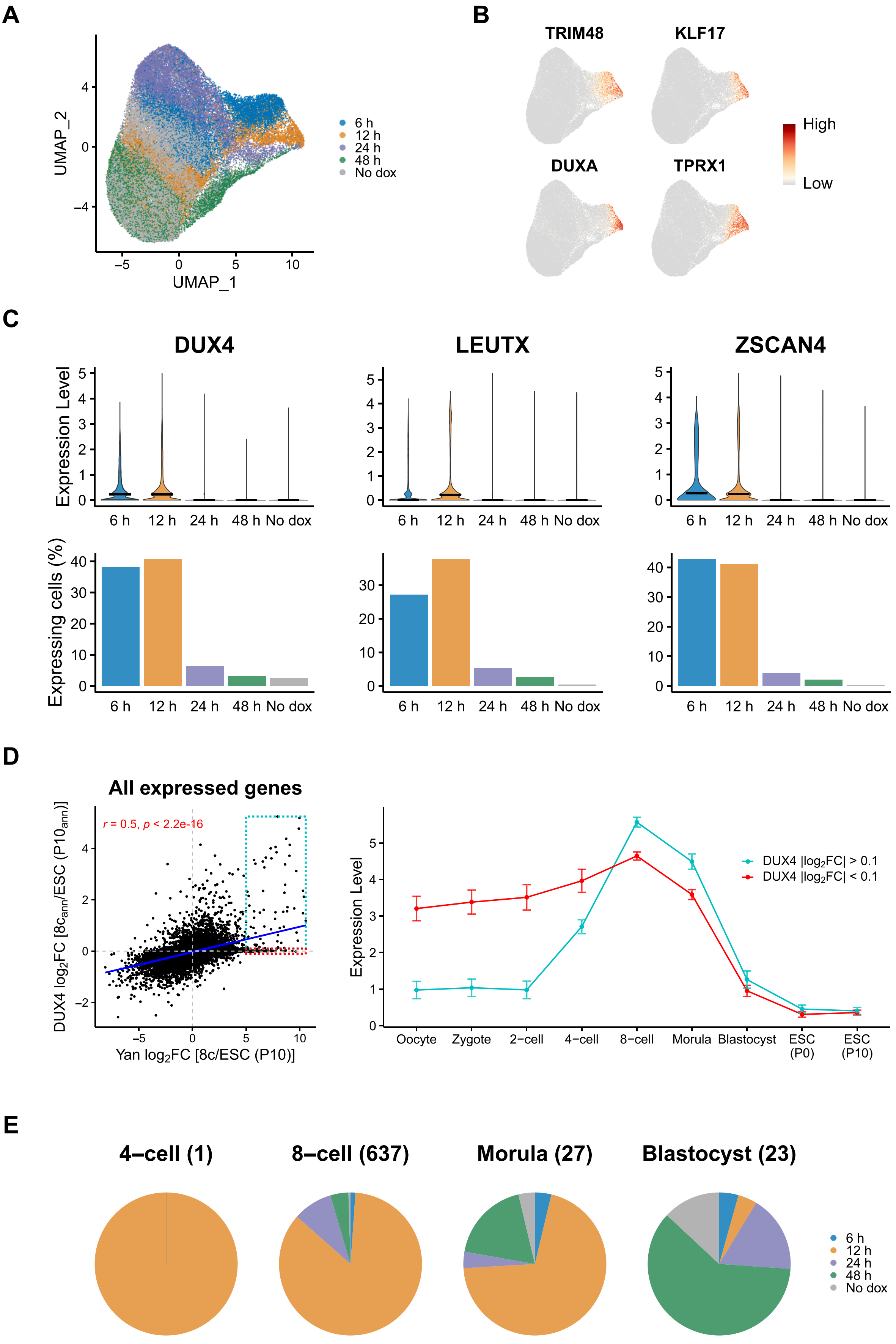

A
UMAP_2
UMAP_1
6 h
12 h
24 h
48 h
No dox
B
TRIM48
KLF17
DUXA
TPRX1
High
Low
C
DUX4
LEUTX
ZSCAN4
Expression Level
Expressing cells (%)
6 h 12 h 24 h 48 h No dox
D
All expressed genes
r = 0.5, p < 2.2e-16
DUX4 log2FC [8c_ann/ESC (P10_ann)]
Yan log2FC [8c/ESC (P10)]
Expression Level
DUX4 |log2FC| > 0.1
DUX4 |log2FC| < 0.1
Oocyte
Zygote
2−cell
4−cell
8−cell
Morula
Blastocyst
ESC (P0)
ESC (P10)
E
4−cell (1)
8−cell (637)
Morula (27)
Blastocyst (23)
6 h
12 h
24 h
48 h
No dox

**Figure S2. Time-course analysis of *DUX4*-pulsed hESCs at single-cell level, related to Figure 2**

(A) UMAP plot colored by collected time points.

(B) Expression pattern of DUX4 target genes projected onto the UMAP plot.

(C) Expression of *DUX4* and its target genes by collected time points. Top: expression levels shown as log normalized UMI counts. Bottom: proportion of expressing cells (UMI count > 1).

(D) Transcriptional changes of 15,902 genes in actual 8-cell stage cells (x-axis) and *DUX4*-pulsed cells annotated as 8-cell stage cells (y-axis) compared with hESCs. Axes show the $\log_2$ fold expression changes over hESCs (P10; x-axis) or cells annotated as hESCs (P10; y-axis). $8c_{ann}$, cells annotated as 8-cell stage cells; ESC ($P10_{ann}$), cells annotated as ESC (P10). Right panel shows the mean expression (log FPKM) during early development of 64 differentially expressed genes ($|\log_2 FC| > 0.1$; green) and 72 unchanged genes ($|\log_2 FC| < 0.1$; red) by *DUX4* induction. Error bars denote SEM.

(E) Proportion of collected time points in cells annotated as early embryonic stage cells.

### Fig. S3

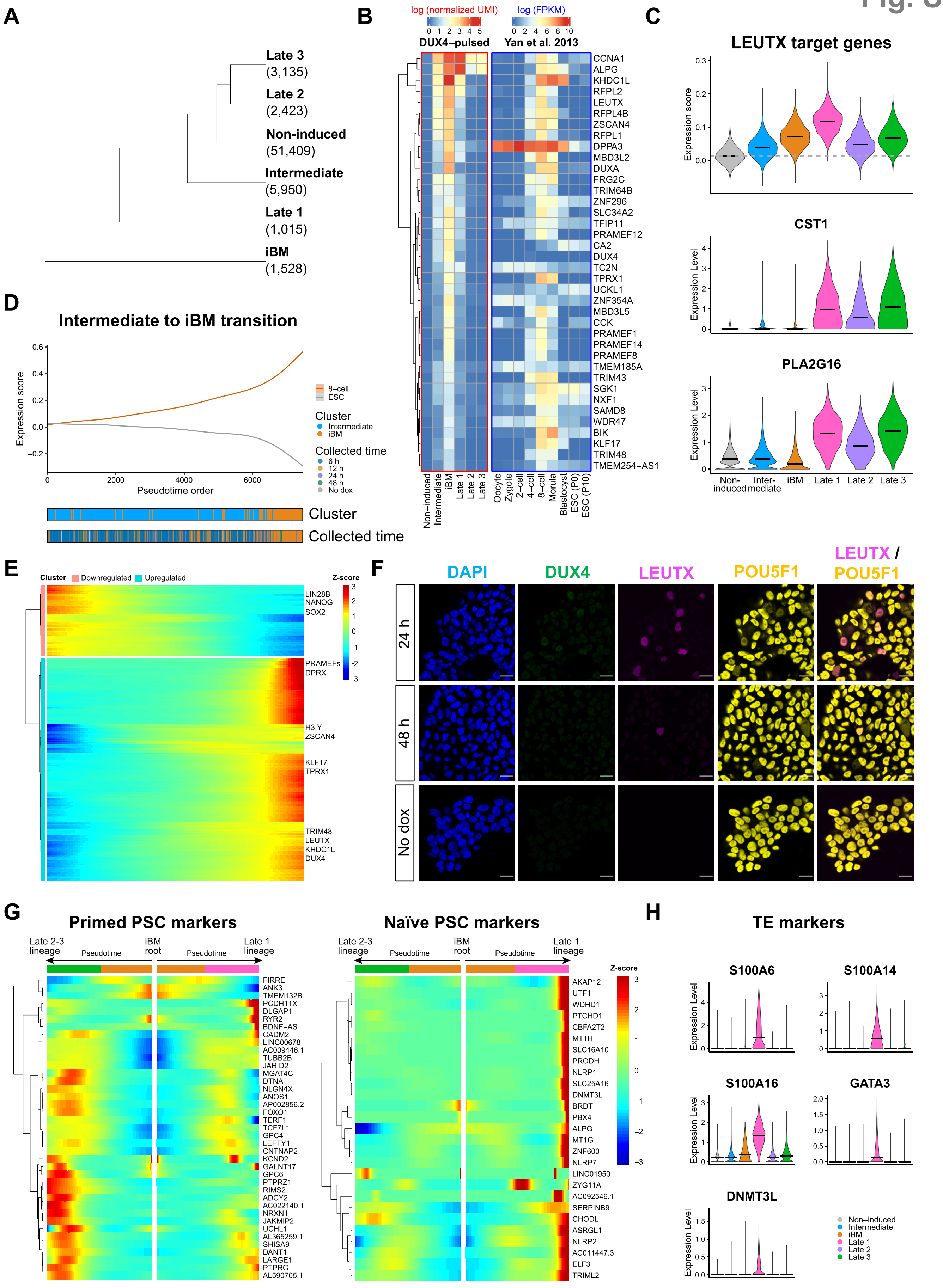
A
Late 3
(3,135)
Late 2
(2,423)
Non-induced
(51,409)
Intermediate
(5,950)
Late 1
(1,015)
iBM
(1,528)
B
log (normalized UMI)
log (FPKM)
DUX4–pulsed
Yan et al. 2013
C
LEUTX target genes
CST1
PLA2G16
D
Intermediate to iBM transition
Pseudotime order
Cluster
Collected time
E
F
DAPI
DUX4
LEUTX
POU5F1
LEUTX / POU5F1
24 h
48 h
No dox
G
Primed PSC markers
Naïve PSC markers
H
TE markers
S100A6
S100A14
S100A16
GATA3
DNMT3L

**Figure S3. Detailed characterization of *DUX4*-pulsed hESCs at single-cell level, related to Figure 3**

(A) Hierarchical clustering analysis of the 6 clusters. Numbers in parentheses indicate the number of cells.

(B) iBM-cluster marker gene expression in *DUX4*-pulsed hESCs in each cluster (left) and human preimplantation embryo (right). Clustering of genes was performed based on the expression pattern across clusters in *DUX4*-pulsed hESCs. See also Table S2.

(C) LEUTX target gene expression in each cluster. LEUTX target gene expression score was calculated with the 299 genes. The horizontal gray dotted line indicates the median score in the Non-induced cluster. *CST1* and *PLA2G16* are representatives. See also Table S3.

(D) Gene expression score changes of 8-cell and ESC in cells from Intermediate and iBM clusters along the pseudotime.

(E) Heatmap of 675 significantly changed genes ($q < 1e-100$) along the pseudotime from Intermediate to iBM transition, clustered by pseudotemporal expression pattern. x-axis corresponds to the pseudotime order shown in Figure S3D. See also Table S4.

(F) Immunocytochemical detection of DUX4, LEUTX, and POU5F1 in *DUX4*-pulsed hESCs at 24 h, 48 h, and without induction (No dox). DAPI (blue) was used as nuclear counterstain. Scale bars, 20 μm.

(G) Expression changes of naïve (left) and primed (right) PSC markers along the pseudotime from iBM to Late transition.

(H) Expression of TE markers in each cluster. The horizontal black bars indicate the median score per cluster.

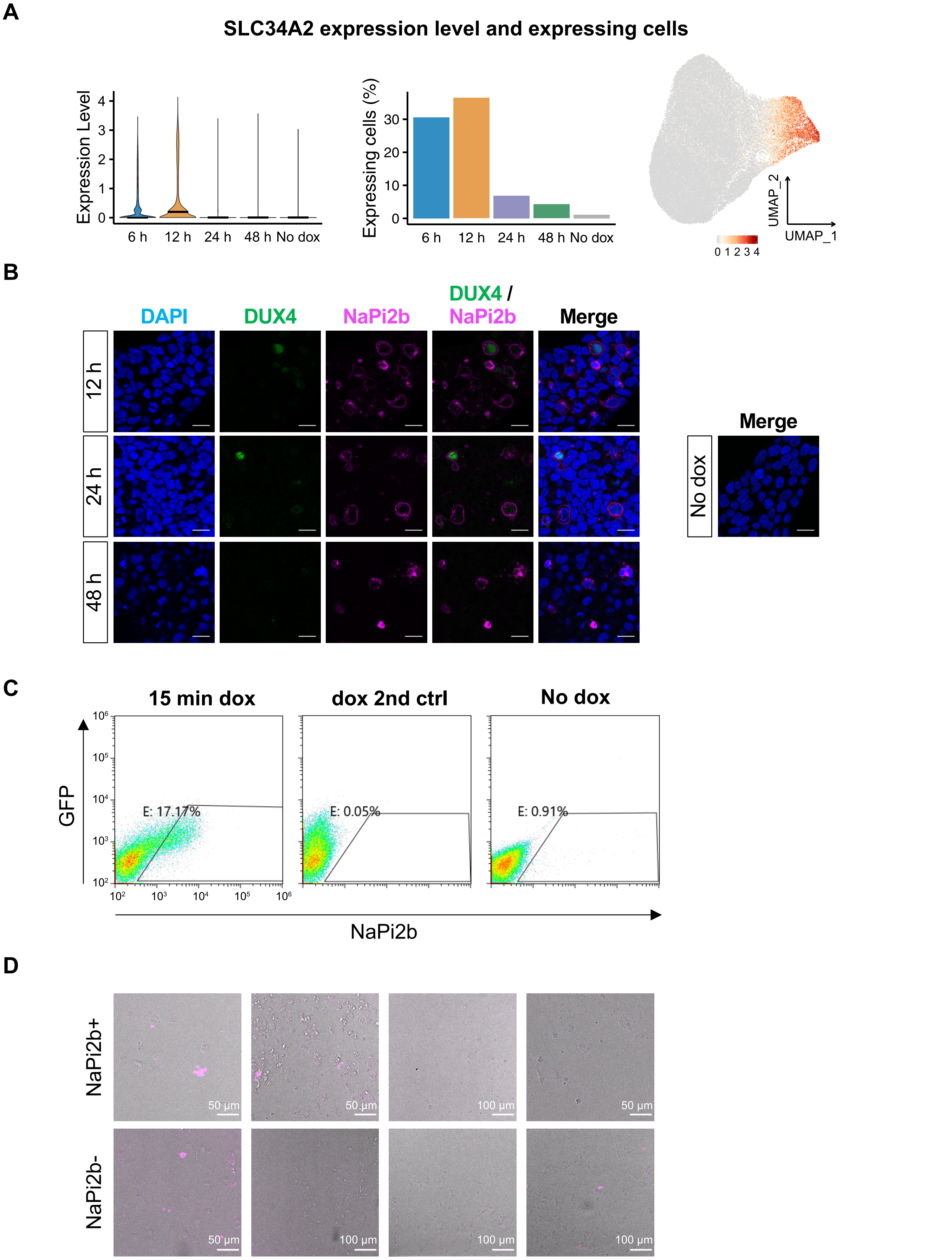
A
SLC34A2 expression level and expressing cells
Expression Level
6 h
12 h
24 h
48 h
No dox
Expressing cells (%)
UMAP_2
UMAP_1
0 1 2 3 4
B
DAPI
DUX4
NaPi2b
DUX4 / NaPi2b
Merge
12 h
24 h
48 h
No dox
C
15 min dox
dox 2nd ctrl
No dox
GFP
NaPi2b
E: 17.17%
E: 0.05%
E: 0.91%
D
NaPi2b+
NaPi2b-
50 μm
100 μm

**Figure S4. iBM cells could be enriched with an anti-NaPi2b antibody, related to Figure 4**
(A) Left: *SLC34A2* expression levels shown as log normalized UMI counts. Middle: proportion of *SLC34A2* expressing cells (UMI count > 1). Right: *SLC34A2* expression projected onto the UMAP plot.
(B) Immunocytochemical detection of DUX4 and NaPi2b in *DUX4*-pulsed hESCs at 12 h, 24 h, and 48 h. Untreated cells (No dox) are shown in right. DAPI (blue) was used as nuclear counterstain. Scale bars, 20 μm.
(C) Flow cytometric analysis showing the gating of NaPi2b-positive cells. Representative data from two independent experiments are shown. 2nd ctrl, secondary antibody control.
(D) Annexin V staining of NaPi2b+/- cells after 6 h of culture. Scale bars are as shown in the images.
